# Activation of the autophagy pathway affects Dengue virus infection in *Aedes aegypti*

**DOI:** 10.1101/2021.04.06.438578

**Authors:** Tse-Yu Chen, Chelsea T. Smartt

**Author notes:** Corresponding author, (C.T.S.).

## Abstract

Mosquito-borne Dengue virus (DENV) has caused major disease worldwide, impacting 50 to 100 million people every year, and is spread by the major mosquito vector *Aedes aegypti*. Understanding mosquito physiology and developing new control strategies becomes an important issue to eliminate DENV. We focused on autophagy, a pathway suggested as having a positive influence on virus replication in humans, as a potential anti-viral target in the mosquito. To understand the role played by autophagy in *Ae. aegypti*, we examined the expression of the pathway *in vitro* (Aag-2 cell) and *in vivo* (*Ae. aegypti*). The results indicated that DENV infection in Aag-2 cells caused the microtubule-associated protein light chain 3-phosphatidylethanolamine conjugate (LC3-II) protein levels to increase which indicated the activation of the autophagy pathway. Rapamycin and 3-Methyladenine were used to activate or suppress the autophagy pathway, respectively. Rapamycin treatment decreased the virus titer in the Aag-2 cells, but the 3-Methyladenine treatment did not affect DENV titer. In *Ae. aegypti*, microinjected rapamycin increased the DENV titer after one-day infection and was significantly different compared to the control group titer. Two ATG genes, ATG4 and ATG12, were expressed differentially under the rapamycin treatments. Although the results differed between *in vitro* and *in vivo* studies, findings from both support the interaction between autophagy and DENV. Our studies revealed the activation of the autophagy pathway through rapamycin could be related to DENV infection in the mosquito. The possibility of autophagy being associated with different antiviral mechanisms at different extrinsic incubation times and tissues in *Ae. aegypti* is discussed.

**Author Summary:** Dengue virus (DENV) has been a great threat to public health and has not developed an efficient method to stop the transmission. To understand the complex interaction between virus and mosquito, we investigate the autophagy pathway and its role during the infection process. We noticed the induction of autophagy pathways from DENV infection in Aag-2 cells and blood meal from *Ae. aegypti*. Moreover, activation of the autophagy pathway from rapamycin could alter the DENV titer. Our results indicated the autophagy pathway is associated with DENV and could be crucial during the DENV infection. Furthermore, we proved the practicality of small molecules in altering the autophagy pathway in mosquitoes, and thus the usage of small molecules as possible mosquito pathogen vaccines should be evaluated.

## Introduction

Dengue virus (DENV) is a single-strand RNA virus, with three structural proteins (capsid protein C, membrane protein M, envelope protein E) and seven nonstructural proteins (NS1, NS2A, NS2B, NS3, NS4A, NS4B, NS5) [1]. Dengue virus is described as having four serotypes, DENV-1, DENV-2, DENV-3, and DENV-4. One disease caused by DENV is dengue fever, and because of the antibody-dependent enhancement, where the antibody against the first serotype cannot neutralize the virus particles of a different serotype [2], some cases will turn into dengue hemorrhagic fever or dengue shock syndrome.

Autophagy is a pathway maintain cellular homeostasis including removal of unnecessary or dysfunctional components in the cell with several autophagy-related proteins (ATG), and controls extensive cellular remodeling [3]. The ATG proteins are required for the process of autophagosome formation. The role of autophagy in cells is to balance sources of energy and in response to stress [4]. Autophagy is also involved in removing misfolded or aggregated proteins [5], clearing damaged organelles [6,7], and eliminating intracellular pathogens [8,9]. Failure in the autophagy function to maintain homeostasis is likely to trigger cell death processes [10]. The involvement of this pathway can be determined by the detection of microtubule-associated protein light chain 3 (LC3) protein. Increased LC3-II levels can be associated with autophagosome synthesis [11,12]. After the maturation of the autophagosome, it fuses with the lysosome to degrade the contents of the autolysosome [13]. Autophagy has also been shown to be involved in immunity where pattern recognition receptors will trigger autophagy to eliminate intracellular microorganisms in humans [14,15] and drosophila [16–18].

Rapamycin is an inhibitor to the mammalian target of rapamycin complex 1 (mTORC1) [19] and therefore, can be used as an inducer of autophagy [20]. On the other hand, 3-Methyladenine (3-MA) is a specific inhibitor of PI3K and is known to interfere with autophagosome formation [21].

DENV infection in humans has been shown to trigger the autophagy pathway [22] and this activation enhances DENV replication [23]. Studies in mammalian cell lines revealed that DENV nonstructural protein NS4A was able to induce autophagy and protected cells from death during infection [24]. Moreover, DENV induced autophagy provided energy resources and released free fatty acids that increase the efficiency of virus replication [25]. Another

DENV nonstructural protein NS1 can also increase LC3-II protein levels as well as cause vascular leakage [26]. The endoplasmic reticulum stress induced by DENV is required for autophagy activation and contributes to viral replication [27]

Although in the drosophila, autophagy has been shown to target intracellular pathogens [28], the interaction between autophagy and virus is not clear in the mosquito. Sindbis virus-infected C6/36 cell activated the PI3K-Akt-TOR pathway, an upstream pathway of autophagy, to enhance cap-dependent translation [29]. In *Ae. aegypti*, after DENV-2 infection, the autophagy-related genes had higher expression levels and the autophagosomes were detected in the midgut [30]. Although the evidence showed the potential interaction between autophagy and DENV, there is still a gap in knowledge about the role of autophagy in the mosquito during viral infections [31].

Here we investigated the role of the autophagy pathway *in vitro* (Aag-2 cell) and *in vivo* (*Ae. aegypti*), using the small molecules rapamycin and 3-MA as activator and inhibitor of the pathway, in DENV infection in mosquitoes. We observed the LC3-II levels increased in the Aag-2 cells after DENV-2 infection. The treatment with rapamycin decreased the viral titer in Aag-2 cells but not under 3-MA treatment. However, we noticed a higher viral titer in *Ae. aegypti* one day after DENV infection from the rapamycin treatment group. Taken together, our results indicate that autophagy might be involved early in the extrinsic incubation period in the mosquito. Further, the autophagy pathway may differ by tissue in mosquitoes.

## Results

### Activation of autophagy in Aag-2 cells and *Ae. aegypti* after DENV infection

To understand the status of autophagy after DENV infection, both *in vitro* and *in vivo* systems were investigated. The Aag-2 cells were infected with DENV-2. After 1 and 2 days, the cell samples were collected and DENV titer measured and analyzed for activation of the autophagy pathway. The western blot showed that LC3-II level was increased on both day 1 and day 2 after infection (Fig 1A) which indicated the activation of the autophagy pathway under DENV infection in Aag-2 cells. Although the DENV-2 triggered autophagy activation *in vitro*, in DENV infected *Ae. aegypti* (4.03±0.53 log (base 10) plaque-forming unit equivalents per milliliter (log PFUe/ml)) the LC3-II was increased 2 and 3 days after blood-feeding in both control and infected groups (Fig 1B).

**Fig 1.**
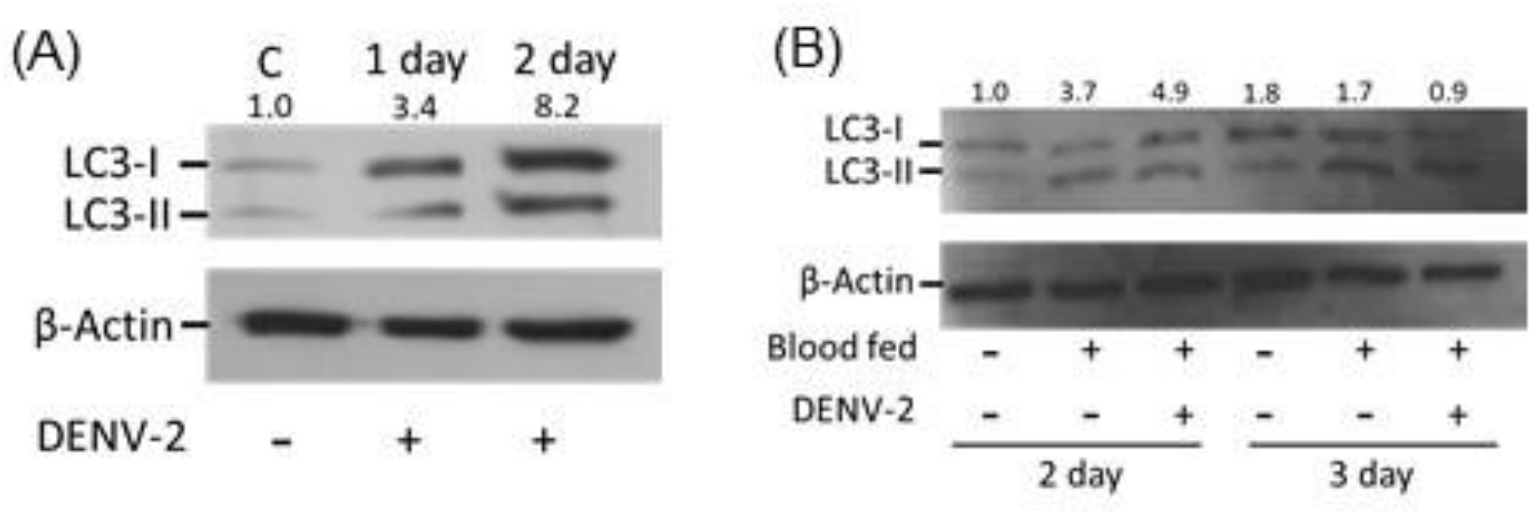
The LC3 protein level in Aag-2 and *Ae. aegypti* after DENV infection. (A) Aag-2 cells were infected with DENV-2 and samples were collected after 1 and 2 days. (B) Five to six day old *Ae. aegypti* were fed with artificial blood with or without DENV-2 and tested for the activation of the autophagy marker LC3-II. β-Actin was detected as internal control. LC3-I = Microtubule-associated protein light chain 3, LC3-II = Microtubule-associated protein light chain 3- phosphatidylethanolamine conjugate, C = Control. The relative ratios of the density were calculated from ImageJ compared between LC3-II and β-Actin are shown.

Several ATG gene expression levels were checked after DENV infection in both *in vitro* and *in vivo* samples. The ATG1 expression in the infection group was increased after 1 day compared to the control group in the Aag-2 cells (S1A Fig). The ATG1 expression level differences, as well as other ATG genes, did not occur after day 2 in DENV infected Aag-2 cells. On the other hand, expression of the ATG genes did not change at any time point tested in DENV infected *Ae. aegypti* compared to blood-fed mosquitoes (S1B Fig).

### Manipulation of the autophagy pathway following exposure with small molecules and DENV infection in Aag-2 cells

Rapamycin was used as an activator of autophagy in the experiment. The 50 nM of rapamycin was applied to Aag-2 cells in culture and protein samples were collected after 1, 3, and 6 hours post-exposure. The LC3-II protein was detected after 1-hour treatment and stably expressed in both 3- and 6-hour samples (Fig 2A). Based on the LC3-II level, 3 hours of rapamycin pre-treatment was chosen to analyze the effects of DENV infection. The virus was incubated in Aag-2 cells for 2 days before being harvested. The LC3-II was induced in the control, pre-treated DMSO groups, and rapamycin pre-treatment group after DENV infection (Fig 2B).

**Fig 2.**
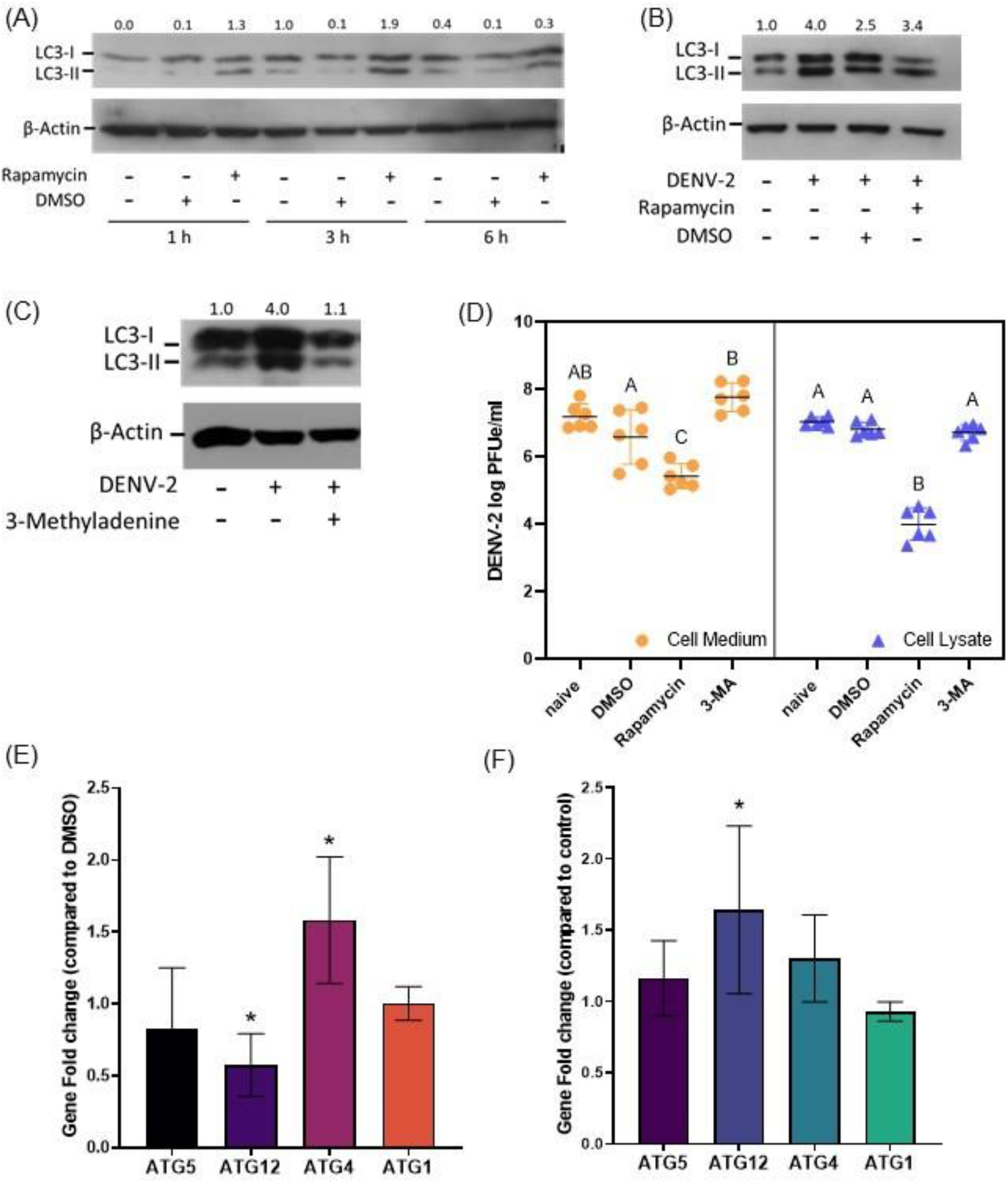
Manipulation of the autophagy pathway with small molecules and after infection with DENV-2 in Aag-2 cells. (A) Western blot results showing LC3-II level after treatment with rapamycin and/or DMSO for 1, 3, and 6 hours. (B) Western blot results showing LC3-II level after rapamycin, DMSO and/or DENV infection. (C) Western blot results showing LC3-II level after treatment with 3-Methyladenine or Schneider’s Drosophila Media and /or infected with DENV. (D) DENV-2 log PFUe/ml titer changes in both Aag-2 cell lysate and cell medium pretreated with DMSO, rapamycin or 3-MA for 3 hours. Holm-Šídák’s multiple comparisons test was used to calculate the *p*-value and levels not connected by the same letter are significantly different. (E) Fold change in ATG transcript levels in cell lysates after pretreatment with rapamycin and DMSO and infected with DENV-2. (F) The ATG transcript level differences in cell lysates after pretreatment with 3-Methyladenine and media and infected with DENV-2. Transcript level differences between control and treatment groups were analyzed using the Wilcoxon Method and *p*-value <0.05 was marked with asterisks (*). 3-MA = 3-Methyladenine, DMSO = dimethyl sulfoxide, ATG = autophagy-related proteins. The relative ratios of the density were calculated from ImageJ compared between LC3-II and β-Actin are shown.

On the other hand, 3-MA was selected as the inhibitor of the autophagy pathway. Three hours after 3-MA pre-treatment of cells, DENV was incubated with Aag-2 cells for 2 days. Previous western blots showed the activation of the autophagy pathway under DENV infection (Fig 2A), but the LC3-II level decreased in the 3-MA treatment group (Fig 2C). The result indicated the autophagy was inhibited by 3-MA in the Aag-2 cells.

After confirmation that both rapamycin and 3-MA function in the Aag-2 cells, both the cells and culture medium were collected, separately, after 2 days of infection. The virus titer was detected in the cell lysate samples; the virus titer in the naïve group was 7.04±0.14 log PFUe/ml, the DMSO treatment group was 6.82±0.20 log PFUe/ml, the rapamycin treatment group was 4.00±0.48 log PFUe/ml and the 3-MA treatment group was 6.72±0.23 log PFUe/ml. The DENV titer between naïve group, DMSO treatment group, and 3-MA treatment groups were not significantly different (*p*-value > 0.05), however, rapamycin-treated cell lysate had significantly lower virus titer than the other groups (naïve vs rapamycin *p*-value = 0.005, DMSO vs rapamycin *p*-value = 0.005, 3-MA vs rapamycin *p*-value = 0.005, Fig 2D). In the cell culture medium samples, the titer had a similar pattern as found in the cell lysates, where the naïve group was 7.20±0.37 log PFUe/ml, the DMSO treatment group was 6.60±0.81 log PFUe/ml, the rapamycin treatment group was 5.43±0.36 log PFUe/ml and 3-MA treatment group was 7.77±0.42 log PFUe/ml. The rapamycin-treated cell medium group contained significantly fewer virions than medium from the other three groups (naïve vs rapamycin *p*-value = 0.005, DMSO vs rapamycin *p*-value = 0.02, 3-MA vs rapamycin *p*-value = 0.005). There was no difference between the cell medium titer of the naïve group and the DMSO treatment group (*p*-value > 0.05), although the 3-MA treatment group had a higher titer than the DMSO group (*p*-value = 0.005). Additionally, the DENV titer of the 3-MA treatment group did not show a significant difference to the naïve group (Fig 2D).

Gene transcription level of four ATG genes was checked after small molecule treatment and 2 days after DENV infection in Aag-2 cells. The related transcripts fold change was calculated compared to the proper control groups. In the rapamycin treatment experiment, the fold change in expression of ATG12 was 0.58±0.21 and was significantly lower than the control (*p*-value = 0.03). Transcription of ATG4 had a 1.58±0.44 gene fold change which was significantly higher than the control (*p*-value = 0.02). Both ATG1 and ATG5 did not show any transcript level differences between rapamycin treatment and control while cells were infected with DENV (Fig 2E). Under the 3-MA treatment, DENV infection in the Aag-2 cells did not alter the transcript levels of ATG1, ATG4, and ATG5. However, ATG12 was detected to have a significantly higher fold change (1.64±0.59, *p*-value = 0.04) compared to the control (Fig 2F).

### Activation of the autophagy pathway through rapamycin influenced DENV titer in *Ae. aegypti*

After confirming the interaction between rapamycin and DENV *in vitro*, the role of autophagy *in vivo* was investigated. Female *Ae. aegypti* were microinjected with different doses of rapamycin, 50 μM, 250 μM, and 500 μM, and collected after 8 hours, 1day, and 2 days. The western blot result showed the autophagy activation was triggered by all doses of rapamycin after 1-day of treatment but 250 μM rapamycin treatment had higher LC3-II levels compared to control and DMSO microinjected mosquitoes (Fig 3A). Two days after microinjection, the LC3-II protein expression remained or increased at all the different doses of rapamycin. Based on the rapamycin microinjection study, 250 μM rapamycin with incubation of 1 day, was used to study effects on DENV.

**Fig 3.**
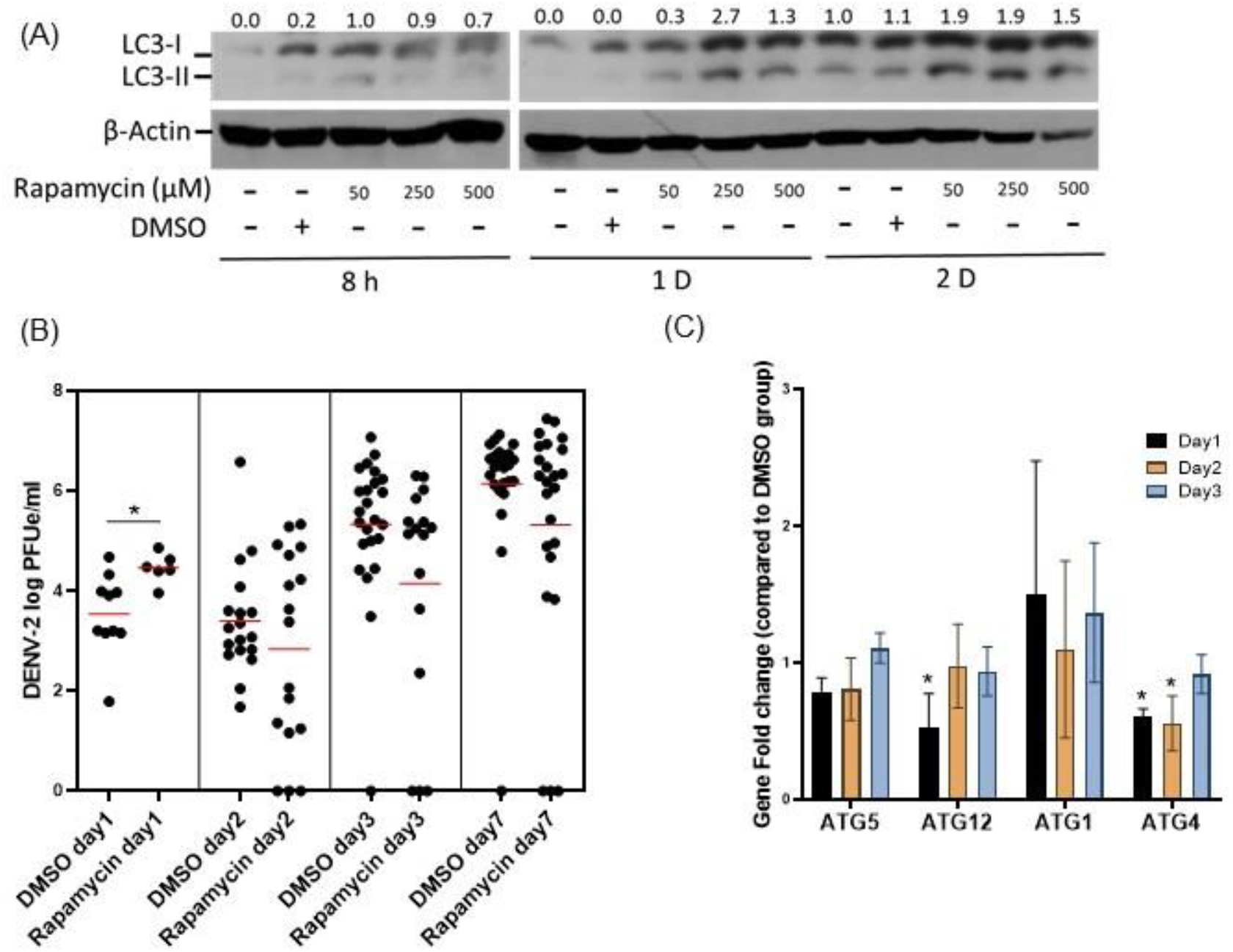
Activation of autophagy pathway through rapamycin influenced DENV replication in *Ae. aegypti*. (A) Western blot results showing LC3-II level in mosquitoes microinjected with DMSO or rapamycin after 8 hours, 1 day, and 2 days. β-Actin was detected as an internal control. (B) DENV-2 log PFUe/ml titer in each mosquito after microinjection with either DMSO or 250 μM rapamycin 1 day before virus infection at 1, 2, 3, and 7 days after infection. (C) The change in ATG transcription in rapamycin injected mosquitoes compared with DMSO injected mosquitoes and infected with DENV-2 over 2 days. The asterisks (*) in 3B and 3C indicates *p*-value < 0.05 between control and treatment groups, calculated by the Wilcoxon Method. LC3-I/II = Microtubule-associated protein light chain 3-I/II, DMSO = dimethyl sulfoxide, ATG = autophagy-related genes. The relative ratios of the density were calculated from ImageJ compared between LC3-II and β-Actin are shown.

The control group (DMSO microinjected) and rapamycin injected group were infected with DENV. The DENV titer of freshly fed control group mosquitoes contained 6.08±0.20 log PFUe/ml and 6.16±0.23 log PFUe/ml in the rapamycin injected group and were not statistically different (*p*-value > 0.05). Samples from both groups were collected after 1, 2, 3, and 7 days after infection and the virus titer evaluated in each mosquito. The average titer on day 1 was 3.54±0.82 log PFUe/ml in the control group (N=10) and 4.46±0.30 log PFUe/ml in the rapamycin injected group (N=6). After 2 days of infection, the DENV titer in mosquitoes of the control group was 3.40±1.16 log PFUe/ml (N=18) and the rapamycin injected group was 2.83±1.97 log PFUe/ml (N=17). In the day 3 samples, the control group contained 5.33±1.43 log PFUe/ml (N=24) and the rapamycin injected group was 4.15±2.28 log PFUe/ml (N=16). After the DENV was incubated in the mosquitoes for 7 days, the virus titers increased to 6.15±1.41 log PFUe/ml (N=24) in the control group and the rapamycin injected group was 5.31±2.29 log PFUe/ml (N=24) (Fig 3B). The results indicated the virus titer was significantly higher under the rapamycin effects after 1-day infection (*p*-value = 0.016), however, the titer in the rapamycin injected group decreased from day 2 and there was no significant difference between the control and rapamycin injected groups on day 2, 3, and 7 (*p*-value > 0.05).

The ATG genes transcripts were examined, and expression was compared under the DENV infection between rapamycin injected mosquitoes and the control mosquitoes. The transcription from four genes (ATG1, 4, 5, and 12) was checked in mosquitoes collected 1, 2, and 3 days after infection. After 1-day infection, the rapamycin injected group had a significantly lower transcription level for ATG4 (0.61±0.06, *p*-value = 0.049) and ATG12 (0.53±0.25, *p*-value = 0.049) but not ATG1 and ATG5 (*p*-value > 0.05). The ATG4 gene was continuously expressed significantly lower in rapamycin injected mosquitoes (0.56±0.20, *p*-value = 0.043). There were no expression differences after day 3 of infection for the four ATG genes (*p*-value > 0.05) (Fig 3C).

## Discussion

Autophagy as a regulated cellular degradative pathway is induced by several factors and has been shown to be associated with DENV infections in humans. The DENV infection triggers the activation of the autophagy pathway and promotes the virus replication [23], however, the interaction between autophagy and DENV in the mosquito is not clear. Here we provided the evidence that activation of the autophagy pathway could interfere with DENV infection in *Ae. aegypti*. The small molecules, rapamycin and 3-MA, were used as activators and inhibitors of the autophagy pathway and showed the activation of the autophagy pathway via rapamycin altered the virus titer *in vitro* and *in vivo*.

The application of the LC3 antibody to detect protein levels of LC3-II is one of the common ways to monitor the activation of autophagy [11,12]. The LC3-II induced by DENV has been shown in several mammalian cell lines, including Huh7 [23], HepG2 [32], and MDCK [24] cells. Here we showed the LC3-II protein level increased in the mosquito cells, Aag-2, after DENV infection (Fig 1A) indicative that the autophagy pathway may be associated with the virus in mosquitoes.

Due to the interaction between virus and cell lines *in vitro*, the difference in LC3-II protein levels in mosquitoes was investigated, including after ingestion of clean blood. We noted the LC3-II protein levels increased after mosquitoes were fed a blood meal (Fig 1B) and these results corroborated what was previously reported [30]. The autophagy pathway is also a key pathway for egg maturation in the mosquito [33] which is strongly associated with the blood-feeding signal. Therefore, it is not surprising that a blood meal without virus is an important signal to trigger the autophagy pathway. The ATG5 gene product is an initiator of autophagosome formation in humans [34] and insects [35], moreover, the DENV replication in humans was shown to be dependent on ATG5 [23]. The ATG5 gene expression level was unchanged after virus infection in both *in vitro* and *in vivo* studies (S1 Fig), however, the results in this study only evaluated between blood fed and virus fed groups. It was previously reported that the ATG5 gene was upregulated following a blood meal [30] and the virus might not affect the transcription level further. We also analyzed expression of several ATG genes shown previously upregulated after blood feeding [30], but only the ATG1 gene transcript level was increased after DENV infection in Aag-2 cells (S1 Fig). The results might suggest that the autophagy-related transcriptomic changes may not be due to DENV infection.

Small molecules, rapamycin, and 3-MA, have been widely used in manipulating the autophagy pathway and understanding the interaction with DENV [22], moreover, both rapamycin and 3-MA were proven useful in studying pathway function in other mosquito systems [36,37]. Therefore, we utilized these molecules in our study (Figs 2A-C and 3A). In the mammalian system, studies have established that DENV activates autophagy upon infection, and uses the autophagic process as part of its replication strategy [23,25,32]. In *Ae. aegypti*, our data indicated the same outcome at early DENV infection time points where the rapamycin injected mosquitoes had higher titers after 1-day infection (Fig 3B), suggesting the interaction between DENV and *Ae. aegypti* might result in activation of autophagy pathway like the mammalian system, at least during the early infection. DENV-induced autophagy is suggested to protect cells against death during infection in mammalian systems [24] and might protect cells in the mosquito, thereby interfering with virus clearance in the mosquito. Surprisingly, activation of the autophagy pathway reduced the virus titer and inhibition of the autophagy pathway slightly increased the DENV titer in Aag-2 cells (Fig 2D). The contradictory results might be influenced by tissue specificity. The *Ae. aegypti* Aag-2 cells are of embryonic origin [38] and are unlike midgut cells, known to be the first tissue contact for the virus in *Ae. aegypti*. This would suggest the interaction between DENV vand the autophagy pathway could be tissue-specific, as seen in mammalian systems. In fact, activation of the autophagy pathway in Aag-2 cells resulted in a significant decrease in virus titer and was similar to a previous mammal monocytic cell study [39]. On the other hand, in *Ae. aegypti*, the activation of the autophagy pathway through rapamycin treatment increased the DENV titer, similar to experiments in mammalian liver cells [23,32]. Further experiments evaluating the effects of DENV infection on tissue-specific activation of autophagy, focusing on the early time points especially during the first 24 hours after infection, should be implemented.

The autophagy pathway has negative feedback that suppresses ATG gene transcription, thus blocking pathway activation [40,41]. Interestingly, two ATG genes, ATG4, and ATG12, had altered transcription levels under rapamycin injection and after DENV infection in both *in vitro* and *in vivo* studies. Both ATG4 and ATG12 are involved in autophagosome formation, and considering their importance in the autophagy pathway, these genes could be potential target genes to investigate as a means to interfere with virus infection or replication in the mosquito.

In summary, we demonstrated the interaction between DENV and the autophagy pathway *in vitro* and *in vivo*. The results indicated the autophagy pathway was a pro-viral mechanism, at early time points post infection in *Ae. aegypti*. However, we noticed the difference between activation of autophagy in the Aag-2 cells compared to in *Ae. aegypti* which could indicate DENV infection interaction with autophagy is different in different mosquito tissues. Given the importance of the autophagy pathway in mosquito physiology including egg maturation, it would be crucial to investigate the influence of autophagy under different extrinsic incubation periods and tissues. Therefore, evaluating the usefulness of the autophagy pathway by gene silencing or small molecules to interfere with virus transmission should be investigated. Moreover, our results showed the practicality of small molecules in altering the autophagy pathway and thus the usage of small molecules as possible mosquito pathogen vaccines should be evaluated.

## Material & Methods

### Mosquitoes

Eggs of *Ae. aegypti* were collected in Vero Beach, Florida in 2015. Mosquitoes were reared at 28°C at 60-80% relative humidity in a climate-controlled room and a light: dark cycle of 14:10 hours. Adults were fed with 5% sucrose solution-soaked cotton rolls. Mosquito colonies were maintained by feeding female mosquitoes on blood from chickens following approved standard protocols (IACUC protocol 201807682) and eggs were collected as described previously [42,43].

### Cell culture

Aag-2 cells were cultured in Schneider’s Drosophila Media (Gibco, Waltham, MA, USA) with 10%FBS and 1% antibiotic-antimycotic (Gibco, Waltham, MA, USA) in a 32°C incubator. African green monkey (Vero) cells were cultured in M199 medium (Gibco, Waltham, MA, USA) in a 37°C incubator with 5% CO2.

### Small molecule and virus infection in Aag-2 cells

Aag-2 cells were seeded into 6-well plates overnight to reach 80% confluency. Rapamycin (Ubiquitin-Proteasome Biotechnologies, Aurora, CO, USA) was dissolved in Dimethyl Sulfoxide (DMSO) (Fisher Scientific, Fair Lawn, NJ, USA) and 3-Methyladenine (Tokyo Chemical Industry Co., Portland, OR, USA) was dissolved in Schneider’s Drosophila Medium. Both small molecules were added into the cell culture directly [44,45]. Each well of Aag-2 cells was treated with rapamycin at 50 nM [23,46] and 5 mM of 3-Methyladenine [23]. Sextuplicate wells for each treatment were made, separately. For both small molecules, cells were pretreated for 3 hours and then infected with DENV-2 at a multiplicity of infection of 0.1 viruses per cell. After the virus was incubated with the cells for 1 hour, the medium was discarded, and the cells were washed with phosphate-buffered saline (PBS, pH 7.4) before fresh medium was added. The infected cells were placed in the incubator for 2 days prior to harvest.

### Small molecule injection of *Ae. aegypti*

Four-day old female *Ae. aegypti* were microinjected with 69 nl of various doses of rapamycin (50, 100, 250 μM) with Nanoject II (Drummond Scientific Co., Broomall, PA, USA). The mosquitoes were collected and tested with LC3 antibody in western blot after 8 hours, 1 day, and 2 days post-injection.

### Infection of *Ae. aegypti*

Previous work showed that 250 μM of rapamycin resulted in the highest level of LC3-II therefore this dose was used for the virus infection study. The injected mosquitoes were transferred to 16 oz cardboard cartons (WebstaurantStore, Lancaster, PA, USA) and placed in an incubator under the same conditions as the climate-controlled room as previously mentioned. One day after microinjection, both the control group (DMSO injected) and the experimental group (rapamycin injected) were fed defibrinated bovine blood (Hemostat, Dixon, CA, USA) containing DENV-2. To prepare the fresh viral media for *Ae. aegypti* infection, the DENV-2 strain New Guinea C (GenBank Accession # KM204118) was inoculated into Vero cells at a multiplicity of infection of 0.01 viruses per cell and incubated at 37°C and 5%CO2 for 5 days. The blood-virus mix was fed to *Ae. aegypti* by use of an artificial feeding apparatus (Hemotek, Lancashire, United Kingdom) with hog casing membranes at 28°C. After a one hour feeding period, fully blood-engorged mosquitoes were collected into cartons and provided 5% sucrose solution and placed at 28°C in an incubator (60-80% relative humidity, 14:10 hours light/dark cycle) for the duration of the experiment. At each time point, mosquitoes were collected and placed at −80°C prior to RNA or protein extraction [42,43].

### Western blot

The cells or mosquito samples were lysed by RIPA buffer (25mM Tris-HCl, 150mM NaCl, 1% NP-40, 1% Sodium deoxycholate, and 0.1% SDS). The samples were incubated with RIPA buffer on ice for 15 minutes and centrifuged at 12000 rpm (13400 g) for 15 minutes. The protein concentration was measured by the BCA protein assay kit (Thermo Fisher, Waltham, MA, USA). The SDS protein sample buffer (MilliporeSigma, Burlington, MA, USA) was mixed with the 35ng protein sample in a 4:1 ratio and then heated at 95°C for 5 minutes. An SDS-PAGE gel (10%) with running buffer (3.03g tris, 14.41 g glycine, and 1g SDS in 1 liter of ddH2O) was used to separate proteins by molecular mass. Proteins in the gel were transferred to nitrocellulose membranes (Invitrogen, Carlsbad, CA, USA) using the transfer buffer (3g tris, 28.8g glycine, and 100 ml methanol in 1 liter of ddH2O) in an Xcell surelock electrophoresis cell (Thermo Fisher, Waltham, MA, USA) under 30 volts for 1 hour. The anti-LC3 antibody was kindly provided by Dr. Alexander Raikhel, Department of Entomology, University of California, Riverside [33], and used as a primary antibody (1:1000) and anti-rabbit antibody (Abcam, Cambridge, MA) as a secondary antibody (1:5000). β-Actin antibody (Cell Signaling Technology, Danvers, MA) was used as the control (1:1000). Clarity Max Western ECL substrate (Bio-Rad Laboratories, Hercules, CA) was applied to stimulate the enzyme horseradish peroxidase (HRP). Signal was detected on X-Omat film (Carestream, Rochester, NY, USA) and film development was applied with Fixer and Developer reagents (Kodak, Rochester, NY, USA).

### RNA extraction, reverse transcription and real-time polymerase chain reaction

TRIzol Reagent (Invitrogen, Carlsbad, CA, USA) was used to extract RNA from Aag-2 cells or mosquito samples. RNA extraction was completed following well-established protocols [43,47], and extracted RNA was stored at −80°C. The RNA samples were treated with RQ1 RNase-Free DNase (Promega, Madison, WI, USA). M-MLV Reverse Transcriptase (Promega, Madison, WI, USA) was used with oligo dT to form cDNA. Transcript levels were determined by Bio-Rad CFX96™ Real-Time PCR. SsoAdvanced SYBR Green Supermix (Bio-Rad, Hercules, CA, USA) and specific primer sets designed for genes in the autophagy pathway, were used to amplify gene products (S1 Table) [30]. *Ae. aegypti* ribosomal protein S7 gene (GenBank Accession # AY380336) was picked as a control for standardizing transcript levels [48]. For quantifying viral titer, the quantitative RT-PCR standardized with plaque assay was applied with equal amounts of RNA (100 ng), iTaq Universal SYBR Green One-step kit (Bio-Rad, Hercules, CA) with DENV-2 specific primers and following a standard protocol (S1 Table) [42,49]. The standard curves for DENV titer were described previously [43] and the virus titer was presented as means of duplicates in log (base 10) plaque-forming unit equivalents per milliliter (log PFUe/ml) [47,49].

## Statistical analysis

Both gene expression data and viral titers were analyzed by the Wilcoxon Method from JMP Pro (www.jmp.com), to calculate the p-value and determine the presence of significant differences between each sample. The figures were made through the GraphPad Prism 9 (www.graphpad.com).

## Acknowledgments

We appreciate Dr. Alexander Raikhel who kindly provided the antibody and Dr. David W. Severson for the autophagy information. The authors would like to acknowledge the excellent technical assistance provided by Sara Ortiz and Natalie Kendziorski.

